# Controlled iris radiance in a diurnal fish looking at prey

**DOI:** 10.1101/206615

**Authors:** Nico K. Michiels, Victoria C. Seeburger, Nadine Kalb, Melissa G. Meadows, Nils Anthes, Amalia A. Mailli, Colin B. Jack

**Affiliations:** Animal Evolutionary Ecology, Inst. Evolution and Ecology, Dep. Biology, Faculty of Science, University of Tübingen, Auf der Morgenstelle 28, 72076 Tübingen, Germany; Universität Hohenheim, Landesanstalt für Bienenkunde (730), August-von-Hartmann-Straβe 13, 70599 Hohenheim, Germany; Lernzentrum Naturwissenschaften, Auf der Morgenstelle 24, 72076 Tübingen, Germany; Biology Department, St. Francis University, 117 Evergreen Drive, Loretto, PA 15940, USA; Nord University, Universitetsaléen 11, 8049 Bodø, Norway; 6 Newcombe Court, Oxford OX2 7NR, UK

**Keywords:** Marine fishes, visual ecology, active sensing, prey detection

## Abstract

Active sensing using light, or active photolocation, is only known from deep sea and nocturnal fish with chemiluminescent “search” lights. Bright irides in diurnal fish species have recently been proposed as a potential analogue. Here, we contribute to this discussion by testing whether iris radiance is actively modulated. The focus is on behaviourally controlled iris reflections, called “ocular sparks”. The triplefin *Tripterygion delaisi* can alternate between red and blue ocular sparks, allowing us to test the prediction that spark frequency and hue depend on background hue and prey presence. In a first experiment, we found that blue ocular sparks were significantly more often “on” against red backgrounds, and red ocular sparks against blue backgrounds, particularly when copepods were present. A second experiment tested whether hungry fish showed more ocular sparks, which was not the case. Again, background hue resulted in differential use of ocular spark types. We conclude that iris radiance through ocular sparks in *T. delaisi* is not a side effect of eye movement, but adaptively modulated in response to the context under which prey are detected. We discuss the possible alternative functions of ocular sparks, including an as yet speculative role in active photolocation.

## 2. Introduction

Some animals enhance their sensory systems by actively sending out signals and perceiving the induced reflections of nearby objects. Well-studied examples of active sensing are echolocation with ultrasound in bats and dolphins, and electrolocation using electric fields in some fishes [1]. “Active photolocation”, where light is used for probing the environment, seems surprisingly rare [1]. Chemiluminescent deep-sea dragonfishes (Fam. Stomiidae), lanternfishes (Fam. Myctophidae) and flashlight fishes (Fam. Anomalopidae) are currently the only vertebrates assumed to use active photolocation [2–8]. They possess a chemiluminescent subocular light organ whose emission facilitates visual detection of prey [9]. The anatomical location of the light organ next to the pupil is optimal to induce reflective eyeshine (bright pupils or “cat’s eyes” effect) in nearby organisms [2]. Reflective eyeshine is a side-effect of the presence of a reflective layer at the back of the eye [10] (ESM 1), which improves vision under dim light [11, 12] or plays a role in camouflage [13]. With such a reflective layer present, focusing eyes in particular backscatter or reflect the incoming light directly back to the source [14]. This can explain why light organs are so close to the pupil in species known to exhibit active photolocation [15].

Interestingly, many diurnal fishes from the euphotic zone show a similar anatomical configuration with conspicuously bright eyes (figure 1, ESM 2). Rather than involving chemiluminescence, light emission originates from reflective and / or fluorescent irides [16–19]. Previously, we found that red fluorescent irides are most prevalent in small fish feeding on small, eyed prey [20], hinting at the possibility that light emission from the iris may constitute a visual aid in detection [21]. Indeed, one of those species captured more prey under “fluorescent-friendly” conditions in a subsequent experiment [22], even though the contribution of active photolocation to this finding remains to be shown.

**Figure 1.**
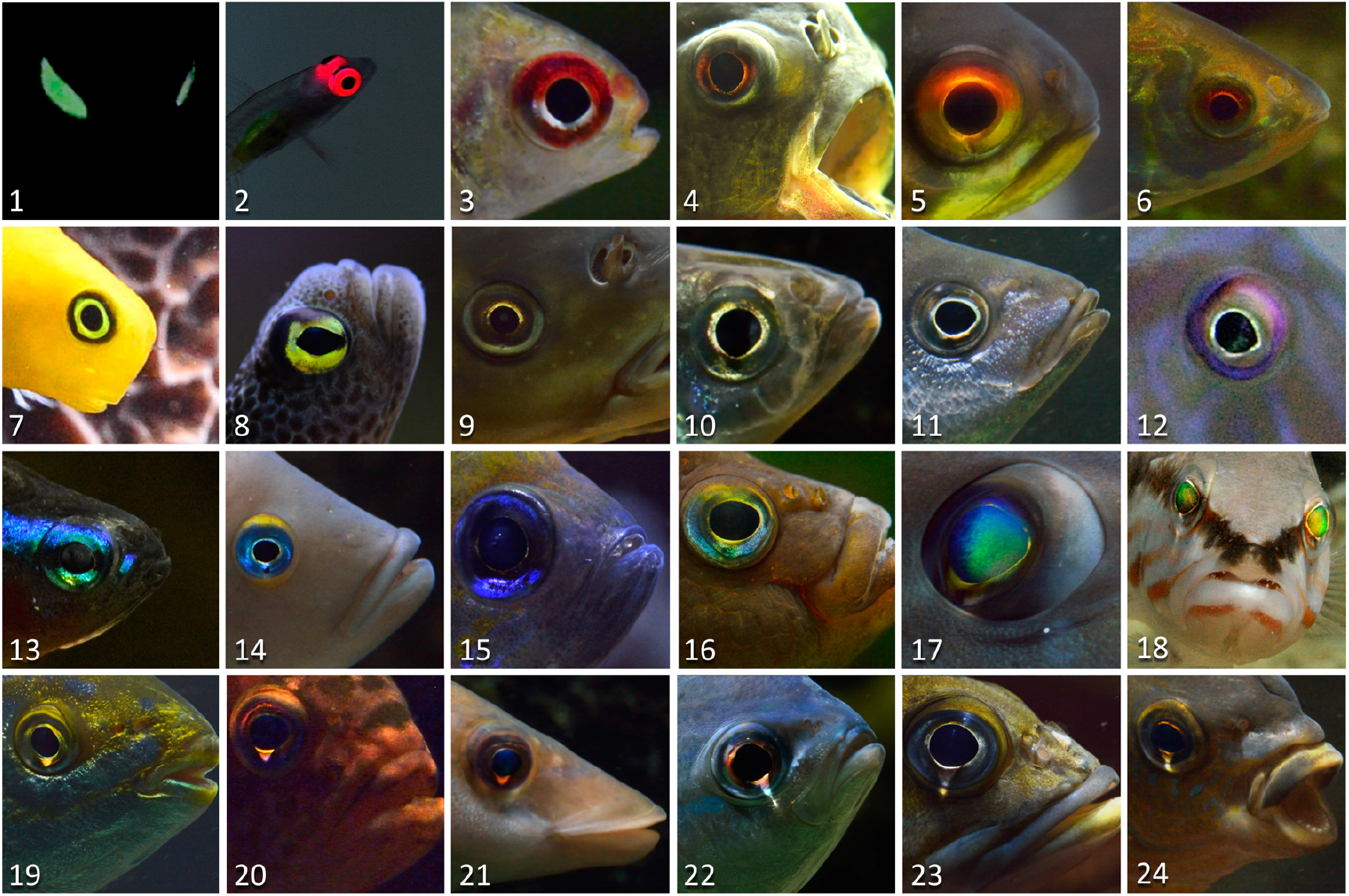
Many fish show bright eyes due to luminescent mechanisms such as chemiluminescence in nocturnal fish (1) or fluorescence in diurnal fish (2). Many more species show forms of reflection off the iris (3-16) or cornea (17-18). This study focuses on so-called “ocular sparks”, a mechanism where fish focus downwelling light onto their own iris (19-22, see figure 2) (species names in ESM 2, photos by N.K.M.).

In this study, we scrutinize an important assumption of active photolocation, namely that diurnal fish emit light “actively”. Specifically, we test the hypothesis that light radiated from the iris is up- or down-regulated in response to the environmental context. Since up- and down-regulation of fluorescence is slow [18, 23] and therefore hard to quantify in free-moving individuals, we focus on the *ocular spark*, a bright spot on the iris, just below the pupil (figure 1, 19-24). Ocular sparks are produced by the protruding spherical lens (figure 2a) which allows downwelling light to cross without entering the pupil, a feature of most bony fish species. It results in a bright focal spot on the lower iris or the skin below the eye. Depending on the pigmentation and structural colouration of the iris at that spot, various radiance spectra can be produced (figure 2c, d). Although ocular sparks can be seen across many fish families (figure 1, 19-24, personal observations), this phenomenon has not been previously described in the literature. Ocular sparks represent a discrete behaviour: Small eye movements are sufficient to switch them “on” or “off” in an instant, facilitating their quantification. Our model species *Tripterygion delaisi* can produce blue ocular sparks that are exclusively reflective (figure 2c) and red ocular sparks that are partly reflective, partly fluorescent, depending on depth [19].

**Figure 2.**
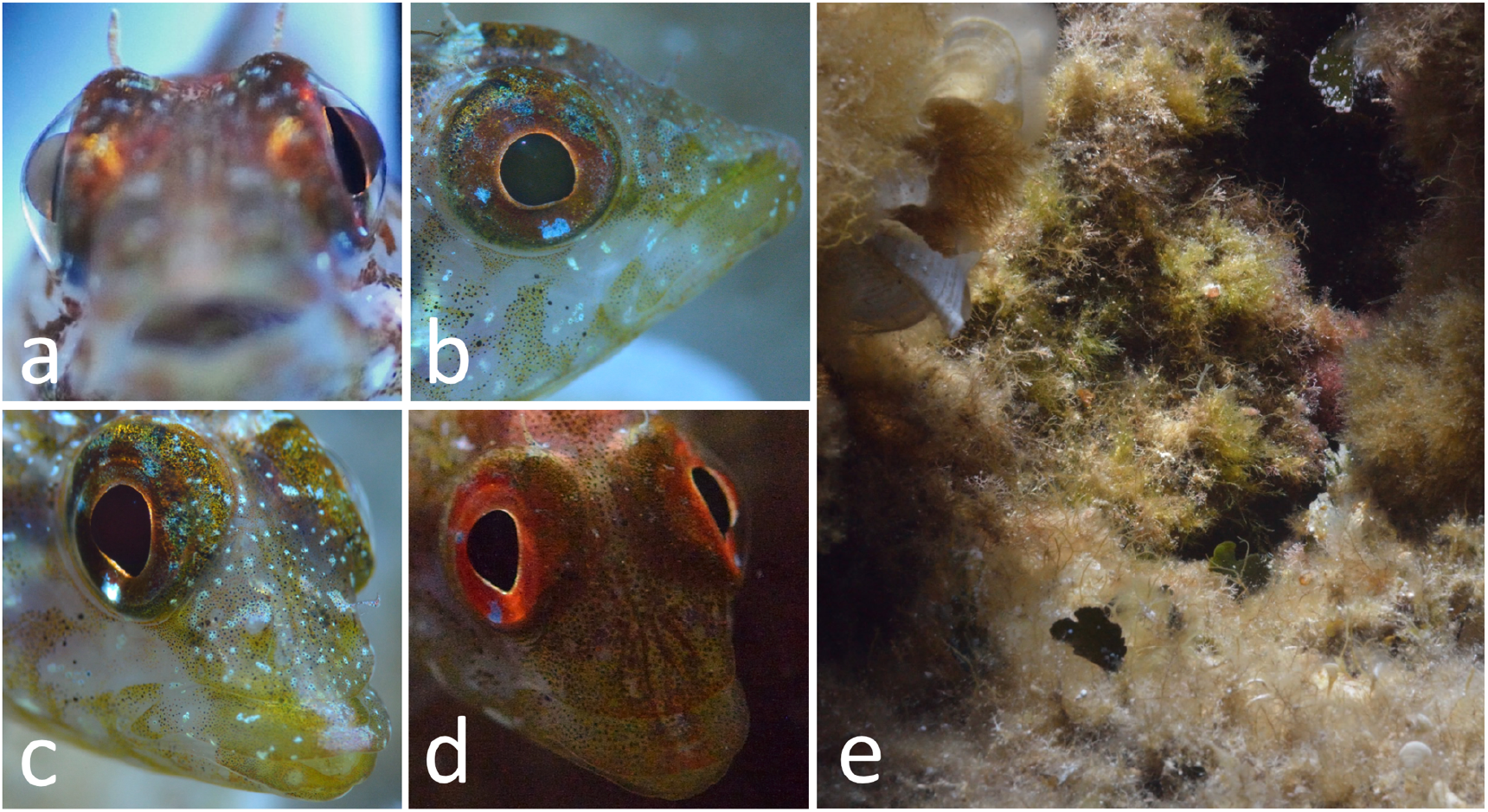
Iris radiance produced by ocular sparks in the triplefin *Tripterygion delaisi*. Downwelling light is focused by the laterally protruding lens of the eye (a, frontal view) onto the lower iris, which features red fluorescent and blue-reflective areas (b). By tilting and rotating the eye, fish can position the focusing spot on a blue iridophore area resulting in a blue ocular spark (c) or a red reflective / fluorescent area, resulting in a red ocular spark (d). (e) A picture of the typical, complex natural environment of *T. delaisi*. See also ESM 3 and 4. All pictures taken in the field by N.K.M.

To test whether the use of ocular sparks correlates with the detection context, we carried out two replicate experiments. In both, we tested whether red ocular sparks are used more when confronted with micro-prey on a blue background, and *vice versa* for blue ocular sparks on red backgrounds. The rationale was that these complementary combinations would maximise the contrast between prey reflection and background. Note that hues of natural substrates at depth can also be red due to fluorescence [16, 24]. The two experiments differed in the second factor tested, being prey presence or absence in the first experiment, and satiation status (hungry or satiated) in the second. We predicted fish to generally show more ocular sparks when prey were present, and satiated fish to be more reluctant to show ocular sparks than hungry fish, assuming a cost of predator attraction when displaying sparks. The results offer support for a differentiated modulation of ocular sparks in response to prey and background hue. We discuss possible alternative explanations for an adaptive use of ocular sparks, including their putative function as a light source in an active photolocation context.

## 3. Material and methods

### 3.1. Model species *Tripterygion delaisi*

The triplefin *Tripterygion delaisi* (Fam. Tripterygiidae) is a 3-5 cm long crypto-benthic fish from the East-Atlantic Ocean and West-Mediterranean Sea. It is common in rocky coastal areas between 5 and 20 m, down to 50 m depth [25, 26], often in or near shady areas and cracks in which they can retreat. During the breeding season (March-May), males have black heads and irides. Otherwise, all age and sex categories show the same cryptic colouration with bright red irides throughout the year. In addition to being fluorescent [19, 27], the iris of *T. delaisi* also features 1-3 white-blue iridophore spots, the biggest one at the ventral edge of the pupil (figure 2b). On these spots, *T. delaisi* can produce a blue, reflective ocular spark (figure 2c). When the focal point of the ocular spark is next to the blue spot, the ocular spark is conspicuously red, which is primarily due to reflection under broad ambient spectra or fluorescence under bluish, deep water conditions [19]. *T. delaisi* sits motionless much of the time while scanning its heterogeneous environment (figure 2e), interrupted by occasional forward hops over short distances (1-10 cm). Free swimming is rare. Potential prey is usually assessed for a few seconds, slowly approached, and then suddenly caught with a sudden strike over 1-3 cm. Preferred prey items are small crustaceans, including harpacticoid copepods [28]. *T. delaisi* moves its eyes independently and with high amplitude, similar – but not as extreme – to the sand lance *Limnichthyes fasciatus* [29]. While doing so, it frequently alternates between both ocular sparks and the “off” state by eye rotation and tilting (ESM 3, ESM 4). Both ocular sparks have been observed in the field down to 30 m. Previous work already indicated that *T. delaisi* can perceive its own fluorescence [19, 30]. The blue ocular spark also falls well within its spectral sensitivity (LWS cone lambda_max_ = 468 nm [19]). Initial prey detection is mostly monocular, but actual striking is often binocular (personal observations by Ulrike Harant and N.K.M.).

### 3.2. Fish collection and housing

We collected *T. delaisi* either near the Station de Recherches Sous-marines et Océanographiques (STARESO) near Calvi, Corsica (France), or near the Hydra Centro Marino Elba in Italy (www.hydra-institute.com). Sampling took place under the general sampling permit of these stations. *T. delaisi* is a non-threatened, non-protected, non-commercial, common species. Animals were transported individually in plastic fish bags filled with 50-100 ml purified seawater and pure oxygen. At the University of Tübingen, fish were kept in individual tanks (*L* x *W* x *H* = 24 x 35 x 39 cm^3^). Black PVC dividers prevented visual contact between neighbouring fish. Coral sand and a piece of rock covered the bottom. All aquaria were interconnected to a flow-through filtering and UV-sterilisation system. The water was kept at a salinity of 35 ‰ at about 23°C. The room used for this study had 21 tanks and was illuminated by weak, diffuse, blue fluorescent ceiling lights (Osram L18W167 Lumilux Blue) for 10 h (0800-1800) per day. From 0900 till 1400 each day, illumination was boosted with one blue LED point source per tank (LED-Fluter AV-LFL10B, Avaloid GmbH, Düsseldorf) (ESM 5). Water quality was checked on a weekly basis. Fish were fed with a mixture of Tetramin (Hauptfutter für alle Zierfische, Tetra GmbH) and Mysis (Einzelfuttermittel, Aki Frost GmbH) every day.

### 3.3. Copepod prey

As experimental prey target, we used *Tigriopus californicus*, an intertidal, benthic, harpacticoid copepod (Crustacea) from the North-American Pacific coast [31]. It is characterised by reflective ocelli (ESM 6, 7). We acquired live samples from http://www.hippozoo.de and cultured them in a climate-controlled room in 20 12 L glass aquaria (30 x 20 x 20 cm3) with artificial seawater (Preis Aquaristik Meersalz) at a salinity of 35 ‰ and 22 °C, with light on 0700-1900 each day. Cultures were fed live phytoplankton daily.

### 3.4. Estimating ocular spark radiance

We measured ocular sparks under controlled conditions in a separate set of individuals in a small tank (L x W x H = 10 x 6 x 4 cm^3^) with a thin (1 mm) glass front in a dark room, illuminated from above. *T. delaisi* remains motionless when placed in an unfamiliar, dark environment. Yet, the fish typically start to investigate their environment using their eyes, thereby turning their ocular sparks on repeatedly. We used a calibrated PR-670 PhotoResearch spectrophotometer with Pritchard optics showing the measurement area as a small black dot in the field of view. We used the smallest dot size of 0.125°. At a distance of ca. 4 cm, it covered about 0.0.25 mm^2^ within the area of the ocular spark. Three measurements were taken from each eye of each fish. Only the brightest measurement of each individual was kept. A Spectralon diffuse white reflectance standard (Labsphere SRS-99-010 AS-01160-060) was placed in a 45° angle in the tank at the position where the fish had been and measured as a reference for the ambient light. Data were transformed into reflectance relative to the diffuse white standard and averaged into a single spectral reflectance curve.

#### Reflectance of blue ocular sparks

To measure blue ocular sparks, we illuminated fish with a liquid light guide attached to a broad-spectrum Leica EL 6000 source. Measurements from one individual included a confounding red fluorescent signal and were discarded, resulting in measurements for 5 fish.

#### Radiance from red ocular sparks

Due to the blue LED illumination of the experimental setup, red ocular sparks were entirely due to fluorescence. Red ocular sparks were therefore measured using a blue Hartenberger Mini Compact LCD dive torch with 7 x 3.5 W 450 nm LEDs for illumination, similar to the monochromatic blue LED illumination used during the experiments (see below). The lens of the spectroradiometer was covered with a LEE 105 Orange filter to suppress the blue excitation light during measurement and improve the signal from the fluorescence emission range. Dividing the measured spectra by the transmission spectrum of the filter yielded the actual radiance spectra shown in the Results. We calculated the relative radiance of the red ocular spark relative to the white standard using the average of the brightest spectra obtained from each of 5 individuals under blue LED light, the relative radiance was subsequently used to estimate the radiance of the red ocular sparks in the experimental setup.

#### Ambient light in the experimental setup

We used a calibrated PhotoResearch PR 740 spectrophotometer to measure the ambient light in the home tanks of the fish as well as the backgrounds against which copepods were presented. In order to assure similar units for all measurements, the downwelling light from the LED source above the aquaria was assessed by measuring the photon radiance of a Spectralon diffuse white reflectance standard and the two types of chambers (red or blue background) in the position where the copepod chamber was presented in the tank (see below). We used a LEE 287 “Double C.T. Orange” filter to suppress the blue ambient light in order to improve the signal in the rest of the spectrum. Data were subsequently corrected for the filter as described above.

### 3.5. Experimental setup

#### General design

The differential usage of ocular sparks was tested in a 2 x 2 design and took place in two replicate runs. Copepod background (blue or red) was the first factor. The second factor was copepod presence or absence in replicate experiment 1, and fish in satiated or hungry state in replicate experiment 2.

#### Experiment 1, with copepod presence or absence as second factor

Each of 21 *T. delaisi* individuals from Elba was exposed to all four treatments in a randomised series. The response of fish to a block of 4 treatments was tested once per week for five weeks, seven individuals per day (Mon, Tue, Wed). We recorded data in the morning, followed by daily feeding in the afternoon.

The stimulus presented to the fish consisted of two microscopy slides (7.6 cm x 2.6 cm) glued together with a “U” shaped line of black, non-toxic, silicone (JBL, AquaSil Silicon), generating a flat, transparent chamber (ca. 25 x 20 x 2 mm3) with a 5 mm opening at the top (ESM 5). Chambers were presented inside the tank at the bottom of the front windowpane facing upward in a 45° angle. For treatments with prey, five live copepods were injected into the chamber using a pipette. Copepods occasionally escaped through the small opening into the aquarium, offering fish a small reward and encouraging them to pay attention. A piece of red fluorescent or blue-grey synthetic felt (Folia Paper, Bringmann, Ton in Ton Bastelfilz) was glued on the back of each chamber to generate the two background hues. Several copies of the chambers were produced and rotated throughout the experiment to balance out individual chamber effects.

Fish behaviour was recorded as a series of 100 still photos, one per 2 s for 200 s, using a GoPro Hero Black Edition camera with macro lens (Schneider optics, B+W macro lens +10) and a filter (LEE Filter 105 Orange) to facilitate distinction between blue and red ocular sparks under the dominant blue ambient light. The viewing angle was adjusted so that the copepod chamber was below the lower edge of the video recording. Thus, image analysis was blind for the treatment. Recording was started remotely when the fish entered the “active zone”, an area within 15 cm from the tank front, indicated with a transparent plastic marker. Within this zone, we expected fish to be able to see the copepods inside the chamber. Three individuals never came into the active zone and were excluded, resulting in a final sample size of *n* = 18 for experiment 1.

#### Experiment 2, with fish satiation state as second factor

We repeated experiment 1 using a new set of 21 *T. delaisi* from Corsica. This time, copepods were present in each trial. We replaced the “copepods present/absent” treatment by a satiation treatment: Fish were exposed to one satiated and one hungry phase per week. An individual started either as “satiated” or “hungry”, alternated across aquaria. Normally, we fed *T. delaisi* one scoop of frozen *Mysis* (~3 shrimps) per day, 7 days per week. Fish in the satiated phase, however, received one scoop of *Mysis* at 1000, 1130, 1300 and 1500 on day 1, followed by two more at 0820 and 0850 just before trials on day 2 (= 6 scoops in ~36 h). On the contrary, fish in the hungry phase received nothing during this period, but were fed after trials on day 2 at 1330 and 1500. On day 3, none of the fish were fed. On days 4 and 5 the same feeding scheme as on days 1 and 2 was applied, but with treatments reversed. All fish received one scoop on day 6 and nothing on day 7, followed by day 1 again. This regime was established during pilot trials and showed a clear difference in responsiveness to feeding in satiated (low response) and hungry (high response) individuals (unpublished results).

During experimental trials, a fish was shown two chambers with 5 copepods in random order with either a blue or red fluorescent background. Each fish was tested for each background (red or blue) six times per satiation status (hungry or satiated). After 2 weeks of acclimation to the procedure, we collected data during the next 4 weeks. Photos were taken every 2 s for 300 s resulting in 150 pictures per fish and treatment. As in the first experiment, we evaluated 100 photos starting with the first photo in which the fish entered the active zone. Two individuals entered the active zone in less than 5 of the 16 trials. Both were excluded from the final analysis, resulting in a sample size *n* = 19 for experiment 2.

### 3.6. Data acquisition

Spark type was assessed visually. Given that fish were free-roaming, each in its own aquarium, the variation in fish illumination, distance and orientation relative to the camera was large. Hence, automated scoring using fixed settings was not feasible. We implemented three measures to prevent observer bias:

(1) Following years of observation in the laboratory and in the field, we derived unambiguous definitions for the two ocular sparks based on a combination of colour, shape, position on the iris, and orientation of the fish and its eye. An example of both sparks in the laboratory can be seen in ESM 4.
(2) Prior to image analysis, scoring persons were trained in correctly applying the criteria from (1) using picture series from the laboratory and the field.
(3) Picture analysis was entirely randomised and blind to the treatment. We counted the number of photos showing (i) a blue ocular spark, (ii) a red ocular spark or (iii) neither. Photos where the fish was pointing away from the chamber were ignored.

### 3.7. Statistical analysis

Data were analysed using a Generalized Linear Mixed Effects Model in the lme4 package [32] of R v3.1.1 [33]. The number of photos showing either type of ocular spark over the total number of photos showing the fish in the active zone was modelled as a binomial (yes/no) response variable with logit link. For the first experiment, the fixed component of all initial models contained the main factors background (blue or red) and copepods (present or absent) as well as their interaction. We further included recording day and treatment order per day to correct for sequence effects on spark usage, and sampling depth to account for differences due to the depth of origin of our experimental fish. Models for the second experiment were constructed in an identical fashion, now with background (blue or red) and satiation status (yes or no) as the main factors and recording day, treatment order per day (the sequence in which each fish was exposed to the two backgrounds) and testing order (the sequence in which the different fish were tested on a given day) as the covariates.

In both models, the initial random component contained Individual ID with random slopes over the background and copepod or satiation treatments. This accounts for the repeated measurements per fish and captures variation that arises from differential individual treatment responses [34]. We further included an observation-level random factor to remove overdispersion [35]. We then performed backward model selection on the random and fixed components using the Bayesian Information Criterion (BIC) to find the most parsimonious model in terms of model fit penalized by the number of parameters [36]. We only report reduced final models, and provide proxies for the goodness-of-fit of the fixed component (marginal R^2^) and the complete model (conditional R^2^) [37, 38] as implemented in the piecewiseSEM package for R [39]. We used Wald z-tests to assess the overall significance of fixed effects. To explore the nature of statistical interactions between our main factors we implemented planned contrasts between the two background types and the presence or absence of copepods (experiment 1) or the satiation treatment within each background type (experiment 2).

Responsiveness of single fish under the two different satiation treatments in the second experiment (latency to reach, and time spent in, the active zone) were analysed using a mixed model with satiation treatment as a fixed factor and individual ID as a random intercept to account for the multiple measurements per fish and treatment using JMP v11.2.

## 4. Results

### 4.1. Spectral properties of ocular sparks

Under the experimental conditions, the photon radiance of blue ocular sparks was about 2 times brighter than that of a diffuse white reflectance standard (figure 3). This coincides with observations in the field, where blue ocular sparks are perceived as the brightest, “sparkling” dot on a substrate by a human observer. The photon radiance of the red ocular spark is bimodal (figure 3, right): It reflects some light in the blue range, but fluoresces in the long-wavelength range, strongly exceeding the ambient light in that range (550-700 nm) under blue illumination.

**Figure 3.**
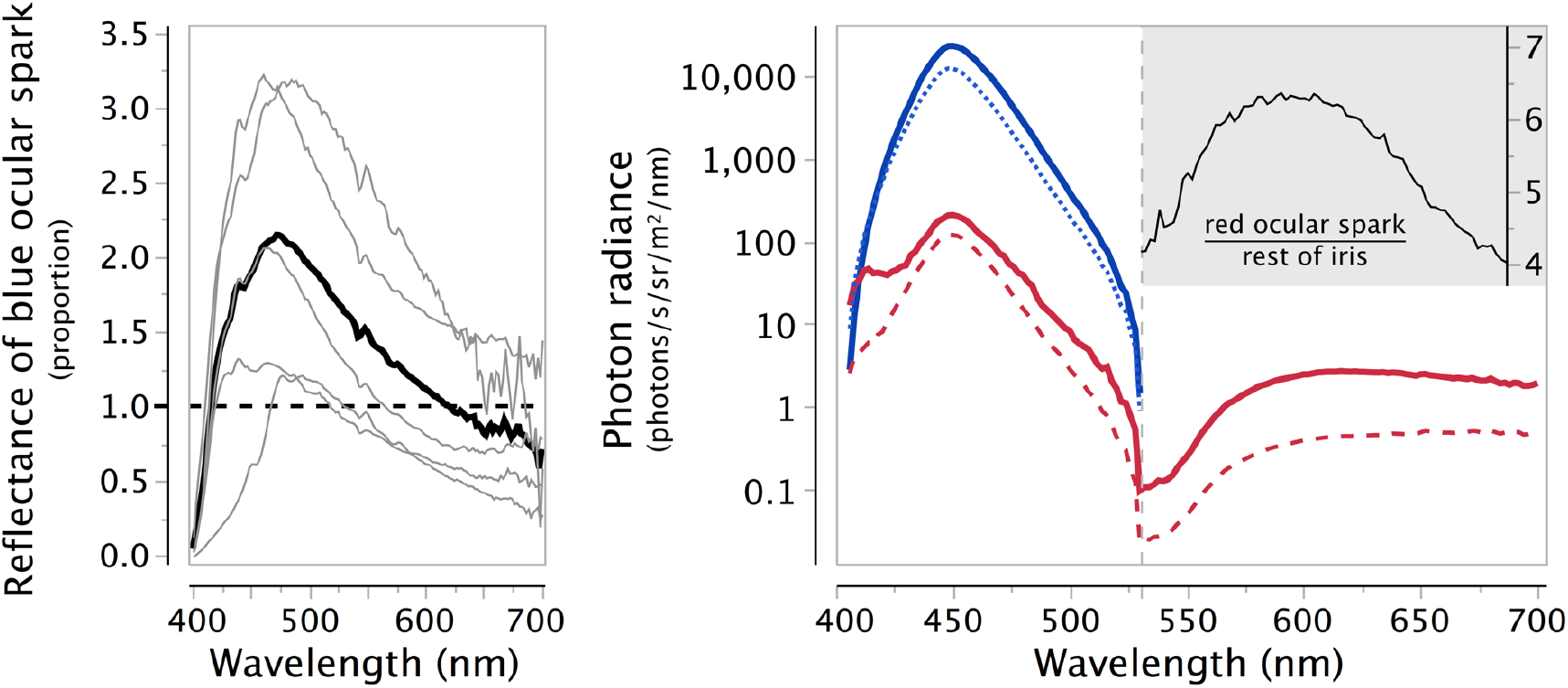
Blue and red ocular spark properties. **Left:** Reflectance (prop.) of the blue ocular spark relative to a white diffuse reflectance standard (dashed line). Thin lines: reflectance spectra from 5 individuals. Thick line: overall average. Note: Due to fluorescence, the red ocular spark cannot be described with a reflectance curve. **Right:** Log10-transformed mean photon radiance under the blue LED illumination of the experiments as a function of wavelength for red ocular sparks (solid red line), other parts of the red fluorescent iris (dashed red line), blue ocular sparks, and a diffuse white reflectance standard (blue dotted line). The insert (top right) shows the brightness of the red ocular spark relative to the fluorescent iris. The vertical dashed line at 530 nm separates the blue LED excitation range and associated reflections (left) from the fluorescent emission range (right).

### 4.2. Experiment 1: Effects of background hue and prey presence

When confronted with copepods in front of a red or blue background, individuals produced blue and red ocular sparks non-randomly (figure 4, table 1): Blue ocular sparks were shown in a significantly higher proportion of pictures against a red background, and red ocular sparks were seen in a higher proportion of pictures against a blue background. The effect weakened, but persisted even in the absence of copepods. Consequently, the interactions between the fixed factors “copepods” (presence/absence) and “background” (red/blue) were also significant.

**Figure 4.**
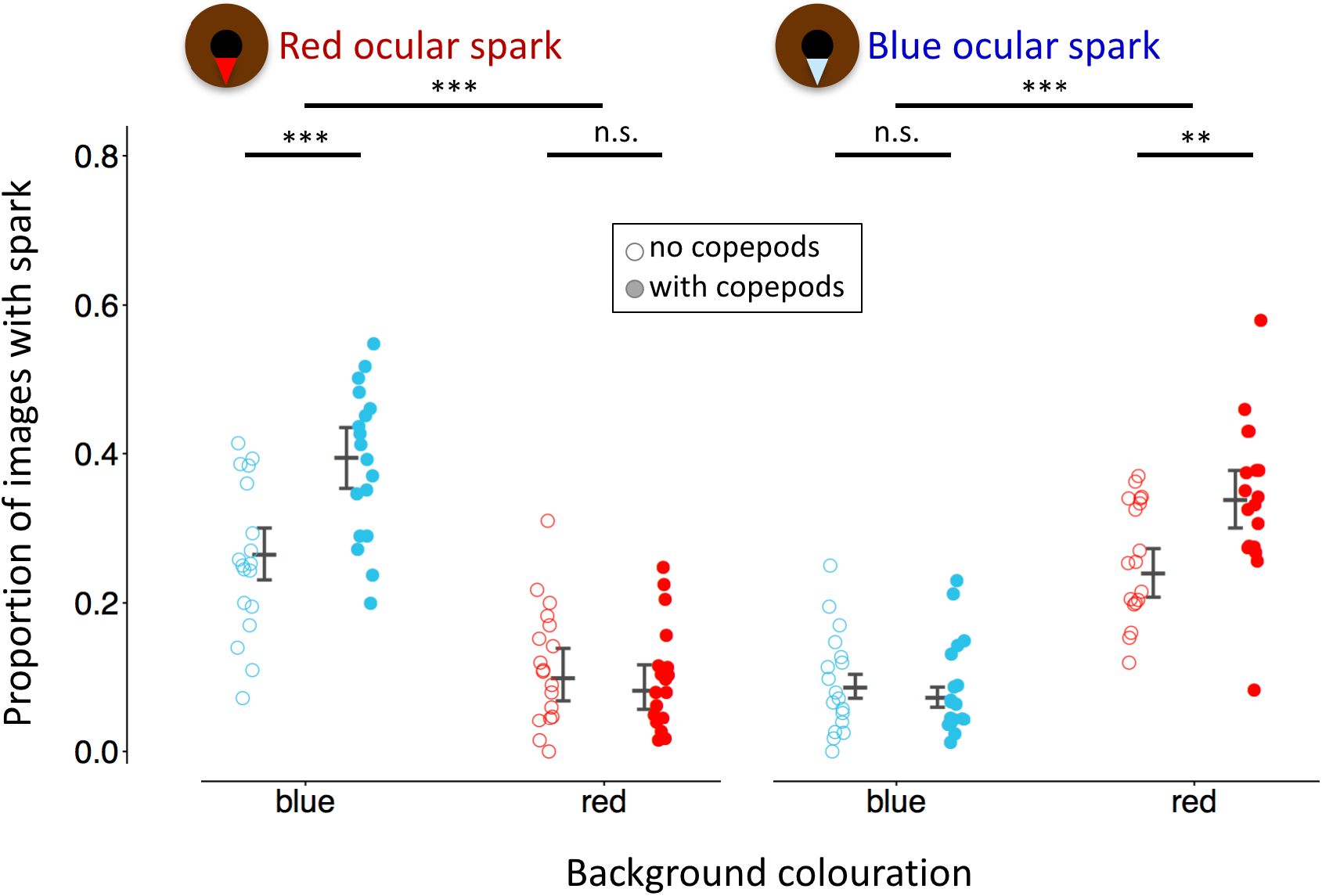
Proportion of observations as a measure of total time during which red (left) and blue (right) ocular sparks were in the “ON” state in the absence (open points) or presence (filled points) of copepod prey against two backgrounds (blue or red). Data points represent averages for each of 18 individuals. Vertical error bars indicate model prediction (central line) with 95% credibility interval (flags) for each treatment group. Significance labels (n.s. = not significant, ** *P* < 0.01, *** *P* < 0.001) indicate planned contrasts between backgrounds, and between copepod treatments within backgrounds.

**Figure 5.**
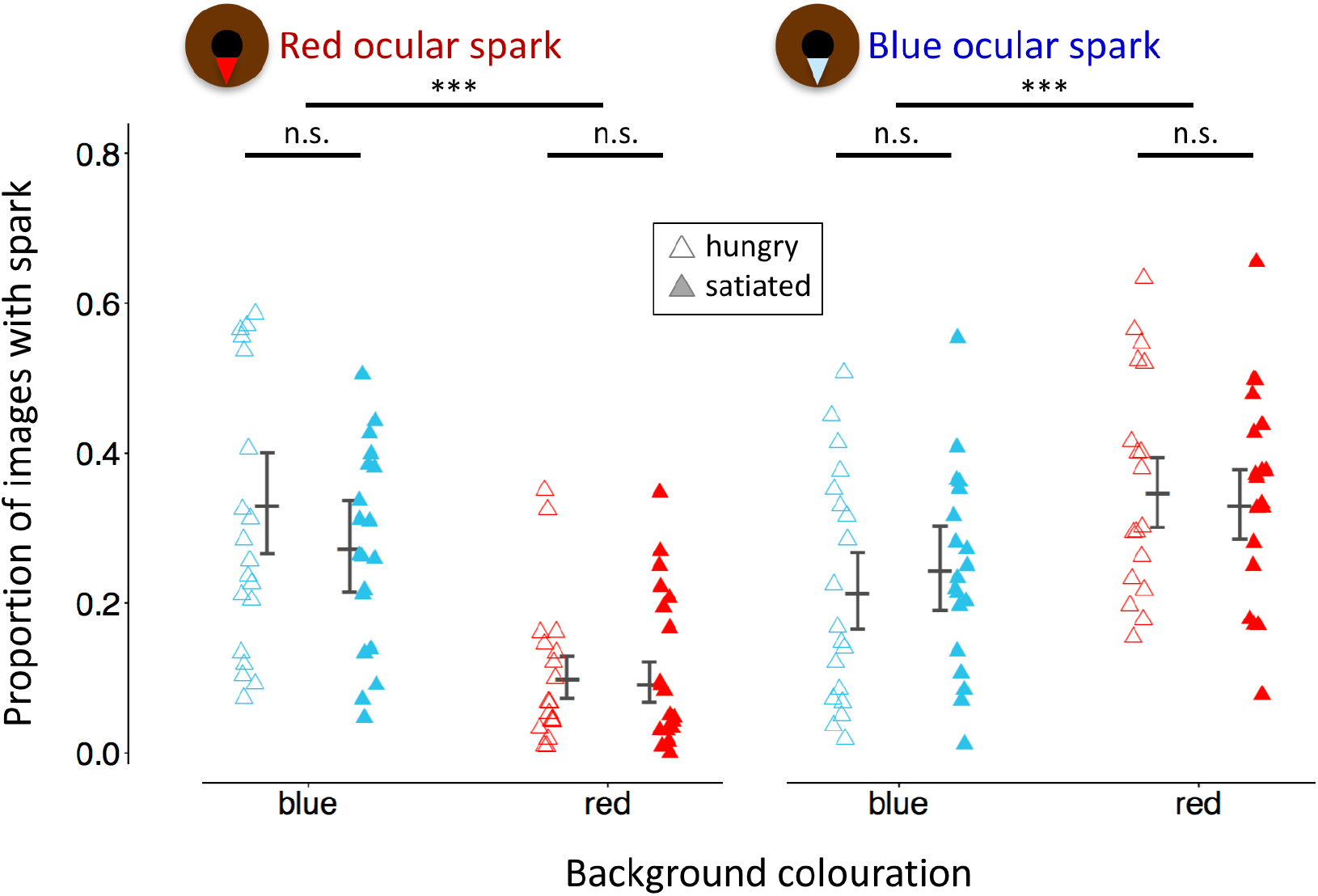
Proportion of observations as a measure of total time during which red (left) and blue (right) ocular sparks were in the “ON” state in hungry (open triangles) and satiated (filled triangles) fish against blue or red backgrounds. Data points represent averages for each of 19 individuals. Vertical error bars indicate model prediction (central line) with 95% credibility interval (flags) for each treatment group. Significance labels (n.s. = not significant, *** *P* < 0.001) indicate planned contrasts between backgrounds, and between satiation treatment within backgrounds.

**Table 1.** Best models from a series of hierarchical GLMM testing the effect of the presence of copepods and background colouration on the total duration (n images) of red (A) and blue (B) ocular sparks in experiment 1 (n = 18 fish).

| **(A) Red ocular spark** |  |  |  |  |
| --- | --- | --- | --- | --- |
| **Parameter** | **Estimate**<br>**(logit-scale)** | **SE** | **$z$** | **$P$** |
| *Intercept* | -1.476 | 0.198 | -7.449 | < 0.0001 |
| *Copepods* | 0.671 | 0.165 | 4.066 | < 0.0001 |
| *Background* | -1.313 | 0.232 | -5.661 | < 0.0001 |
| *Treatment order* | 0.132 | 0.057 | 2.324 | 0.0201 |
| *Copepods\*Background* | -0.987 | 0.255 | -3.830 | 0.0001 |
| **(B) Blue ocular spark** |  |  |  |  |
| **Parameter** | **Estimate**<br>**(logit-scale)** | **SE** | **$z$** | **$P$** |
| *Intercept* | -2.895 | 0.169 | 17.104 | <0.0001 |
| *Copepods* | -0.189 | 0.237 | -0.796 | 0.4263 |
| *Background* | 1.549 | 0.227 | 6.809 | <0.0001 |
| *Copepods\*Background* | 0.721 | 0.316 | 2.281 | 0.0225 |

Marginal R<sup>2</sup>: 17.1 % for red and 17.2 % for blue ocular sparks. The sequence order in which treatments were offered had an effect in (A) and was therefore maintained in the model to compensate for its effect. All other parameters that were initially in the model were removed because of their insignificant contribution (See Methods 2.7). Parameter significance is assessed using Wald $z$ -tests.

### 4.3. Experiment 2: Effects of background hue and satiation status

The replicate experiment again revealed a significant effect of background colour on the use of ocular sparks, similar to that seen in the first experiment (figure 4, table 2). Satiation status, however, showed no detectable effect, irrespective of background hue. The latency with which individuals appeared in the active zone after the start of a trial and the total time spent in the active zone also did not differ between satiation treatments (mixed model satiation treatment effects, *P* = 0.63 and *P* = 0.13).

**Table 2.** Best models from a series of hierarchical GLMM testing the effect of satiation status and background on the total duration *(n* images) of red (A) and blue (B) ocular sparks in experiment 2 (n = 19 fish).

| **(A) Red ocular spark** |  |  |  |  |
| --- | --- | --- | --- | --- |
| **Parameter** | **Estimate**<br>**(logit-scale)** | **SE** | **$z$** | **$P$** |
| *Intercept* | 0.022 | 0.482 | 0.047 | 0.963 |
| *Satiation status* | -0.297 | 0.287 | -1.033 | 0.302 |
| *Background* | -1.963 | 0.308 | -6.369 | <0.0001 |
| *Observation day (1-8)* | -0.147 | 0.047 | -3.124 | 0.002 |
| *Satiation status \* Background* | 0.121 | 0.436 | 0.277 | 0.781 |
| **(B) Blue ocular spark** |  |  |  |  |
| **Parameter** | **Estimate**<br>**(logit-scale)** | **SE** | **$z$** | **$P$** |
| *Intercept* | -1.319 | 0.247 | -5.325 | <0.0001 |
| *Satiation status* | 0.216 | 0.193 | 1.115 | 0.265 |
| *Background* | 0.808 | 0.235 | 3.439 | 0.0005 |
| *Fish testing order* | -0.026 | 0.012 | -2.173 | 0.0298 |
| *Satiation status \* Background* | -0.223 | 0.271 | -0.824 | 0.4099 |

Marginal R: 13.8 % for red and 3.3 % for blue ocular sparks. Significant confounding factors were kept in the model to compensate for their effect (Variables “observation day rank number” for red ocular sparks and “daily fish testing order” for blue ocular sparks). All other parameters that were initially in the model were removed because of their insignificant contribution (See Methods 2.7). Parameter significance is assessed using Wald $z$ -tests.

## 5. Discussion

This study introduces ocular sparks as a new mechanism of iris radiance in fishes and shows that they are actively modulated in response to prey presence and background hue in a cryptobenthic triplefin. Spark use increased in the presence of prey, which is reminiscent to increased pulse rates in echolocating predators when prey have been detected [40, 41]. Moreover, the complementary adjustment of spark hue to background hue is suggestive of visual contrast optimization. Given that spark hue was also adjusted in the absence of copepods, fish seem to adjust baseline sparking to conditions under which they have seen copepods before. Contrary to our expectations, hungry fish did not show an increase in ocular sparking relative to satiated fish (experiment 2). Although satiated fish had shown reduced interest in standard food in a pilot study, the live copepods presented in experiment 2 may have revived their interest. The results lend some support to the hypothesis that ocular sparks may be used in active photolocation and provide the impetus for further experiments.

### Are ocular sparks too weak?

Despite the fact that the radiance of the ocular sparks clearly exceeds that of a 99% white diffuse standard under the experimental conditions, it is nevertheless quite likely that ocular sparks too weak to enhance regular vision in many natural situations. However, they may offer an advantage under specific conditions. Even a weak light source (e.g. a piece of white paper) can induce perceptible reflections over short distances in highly reflective targets and under specific light gradients (e.g. ESM 7, 8). Hence, a benthic micro-predator such as *T. delaisi* exposed to the sun might be able to induce perceptible reflections in the eyes of cryptic organisms hiding in the shade nearby. The issue is therefore not only whether ocular sparks are strong enough, but also whether potential targets are strongly reflective, detection distances short, and prevailing light conditions favourable. Hence, the question whether ocular sparks function as a light source for active photolocation cannot be rejected or accepted by default, but must be assessed in a comprehensive, context-specific manner.

### Do ocular sparks offer advantages over other forms of iris radiance?

Even when assuming iris radiance is useful, why in the form of an ocular spark? It does not radiate more light than reflection off the full iris could, but merely concentrates it into a small spot. Many fish species have silvery or golden irides that reflect light from most or all of their surface (figure 1). One answer may be that light concentrated onto a small spot is less conspicuous to a predator than a full, bright iris. Seen from a distance, an ocular spark will fall below the spatial acuity of most observers, becoming a point source. Any further increase in distance would result in an exponential decrease in radiance [42]. As a result, ocular sparks may be less conspicuous than a complete iris at distance. This may allow micro-predators to hunt successfully over very short ranges, while remaining cryptic over the detection distances used by their own predators. Second, only ocular sparks allow for accurate and immediate swapping between two hues with minimal eye movement. This appears particularly useful for fish foraging on complex substrates. Non-cryptic fish feeding against a homogeneous background such as the open water may not need such a “filter wheel”. Finally, by focusing light onto the lower edge of the pupil, ocular sparks approximate a fish’s equivalent of an ophthalmoscope. Given that its retinal *area centralis* can be positioned in line with this bright, reflective edge and the target (unpublished data), this coaxial arrangement is ideal to highlight eyeshine [2, 21, 43].

### Alternative functions of ocular sparks

As a non-exclusive alternative explanation, ocular sparks might act as lures and *attract* copepods as has been proposed for the fluorescent tentacles of hydromedusae that are assumed to attract fish [44, 45]. This however, seems unlikely for prey with poor visual acuity as is true for benthic copepods with just three ocelli [31]. It also does not explain why fish would alternate between two hues in response to the background of the copepod rather than that of the fish itself. Moreover, *T. delaisi* does not lure or ambush approaching prey, but actively searches for prey. Finally, zooplankton shows an escape response when confronted with flashes of light [46, 47]. Hence, although light-emitting lures exist, it does seem to be a likely function for ocular sparks.

A more plausible alternative function is that ocular sparks are used for intra-specific signalling, as already suggested for the chemiluminescent flashes of *Anomalops katroptron* [2]. *T. delaisi* usually forms loose groups of 2-6 adults and juveniles. It is likely that they can see each other’s ocular sparks. Most behavioural interactions between individuals are, however, characterised by fin-related behaviour such as dorsal fin flicking, push-ups and tail-flicks [48]. Whether ocular sparks supplement this repertoire is as yet unknown, but even if this were the case, it cannot explain ocular spark modulation in solitary individuals in response to prey and background hue.

## 6. Conclusion

The triplefin *T. delaisi* redirects light from its irides in a controlled way by means of ocular sparks in response to prey presence and background hue. While these findings clearly show differential use of ocular sparks, they do not allow to conclude that the fish in the experiment induced and perceived reflections in the copepods. This “detection component” of active photolocation is subject to ongoing experiments and visual modelling.

## Acknowledgments

We are grateful to the many students who contributed to optimising the methodology required for this study: Maximillian Statler, Dorothea Möhrle, Tim Piel, Lisa Kettemer, Johanna Werminghausen and Hannes Nau. We sincerely thank our technical staff for their technical support (Gregor Schulte), fish maintenance (Oeli Oelkurg, Christopher Rader and Martina Hohloch) and copepod breeding (Martina Hohloch, Christopher Rader). This work greatly benefited from comments on previous drafts and discussions from six anonymous reviewers, Pierre-Paul Bitton, Connor M. Champ, Roland Fritsch, Matteo Santon, Magnus Johnson and Ulrike Harant. We also are indebted to the scientists that attended a workshop on active photolocation in November 2014 in Tübingen: João Coimbra, Sönke Johnsen, Almut Kelber, Daniel Osorio, Shelby Temple and Annette Werner. Their critical and constructive suggestions strongly contributed to the quality of our work and eventually also this paper.

## Ethical statement

For the experiments, animals had been well habituated to their “home” aquaria for 6 months. To prevent fish from being distracted by increased human activity in the room, we covered the front of all tanks with a black PVC panel for the duration of the study. Animal husbandry was in accordance with German animal welfare legislation (Tierschutzgesetz of 7 August 2013, §2 and §11) and is checked regularly by independent, regional authorities (Dr. S. Wheeler, Landkreis Tübingen, Veterinärwesen und Lebensmittel-Überwachung, Wilhelm-Keil-Str. 50, 72072 Tübingen, Germany). Data collection relied exclusively on the spontaneous curiosity of fish for the stimulus provided in their home aquarium. Fish were never handled in any way. Consequently, our design did not involve pain, suffering or injury to the animals. According to §7 (2) of the German animal welfare act (Tierschutzgesetz) of 7 August 2013 this implies that a formal application for an ethics permit is not required, as approved by the institutional animal welfare officer, Dr. Annette Denzinger (Fachbereich Biologie, Universität Tübingen, Auf der Morgenstelle 28, 72076 Tübingen, Germany).

## Funding statement

We acknowledge generous support by the German Science Foundation Koselleck grant (Mi482/13-1) and the Volkswagen Foundation (Az. 89148 and Az. 91816) to N.K.M. as well as running support by the University of Tübingen.

## Data accessibility

Data for spectral properties of ocular sparks, copepod eye reflectance and all data of the two experiments are available at DOI: https://doi.org/10.5061/dryad.6cp6f [49]

## Competing interests

The authors declare no competing interests.

## Authors’ Contributions

The idea for active photolocation was brought to us by C.B.J. in 2009. Concept and experimental design were developed by N.K.M. and M.G.M. with assistance from V.S., N.K. and N.A. Spectrometry was developed during student projects under supervision of N.K.M. and M.G.M. and carried out by N.K.M. and A.M. for the copepods (ESM 6). Behavioural experiments were carried out by V.S. (experiment 1) and N.K. (Experiment 2). Data analysis was done by N.A. and N.K.M. The paper was written by N.K.M.

All authors approved the final version of the manuscript.

